# Triton-X 100-treated virus-based ELLA demonstrates discordant antigenic evolution of influenza B virus haemagglutinin and neuraminidase

**DOI:** 10.1101/2024.07.09.602673

**Authors:** Thi H. T. Do, Michelle Wille, Adam K. Wheatley, Marios Koutsakos

## Abstract

Neuraminidase (NA)-specific antibodies have been associated with protection against influenza and thus NA is considered a promising target for next-generation vaccines against influenza A (IAV) and B viruses (IBV). NA inhibition (NI) by antibodies is typically assessed using an enzyme-linked lectin assay (ELLA). However, ELLA can be confounded by anti-hemagglutinin (anti-HA) antibodies that block NA by steric hindrance (termed HA interference). While strategies have been employed to overcome HA interference for IAV, similar approaches have not been assessed for IBV. We found HA interference is common in ELLA using IBV, rendering the technique unreliable. Anti-HA antibodies were not completely depleted from sera by HA-expressing cell lines and this approach was of limited utility. In contrast, we find that treatment of virions with Triton-X 100, but not Tween-20 or ether, efficiently separates the HA and NA components and overcomes interference caused by anti-HA antibodies. We also characterise a panel of recombinant IBV NA proteins that further validated the results from Triton-X 100-treated virus-based ELLA. Using these reagents and assays we demonstrate discordant antigenic evolution between IBV NA and HA over the last 80 years. This optimized ELLA protocol will facilitate further in-depth serological surveys of IBV immunity as well as antigenic characterisation of the IBV NA on a larger scale.

**Importance:** Influenza B viruses contribute to annual epidemics and may cause severe disease, especially in children. Consequently, several approaches are being explored to improve vaccine efficacy, including the addition of neuraminidase. Antigen selection and assessment of serological responses will require a reliable serological assay to specifically quantify Neuraminidase inhibition. While such assays have been assessed for influenza A viruses, this has not been done of influenza B viruses. Our study identifies a readily applicable strategy to measure inhibitory activity of neuraminidase-specific antibodies against influenza B virus without interference from anti-hemagglutinin antibodies. This will aid broader serological assessment of influenza B virus-specific antibodies and antigenic characterisation of the influenza B virus neuraminidase.

## Introduction

Influenza B virus (IBV) infections arise in annual epidemics and can comprise up to 80% of the influenza burden in some years, resulting in significant health and social-economic impacts [1–4]. Similar to IAV, there are two major surface glycoproteins, the hemagglutinin (HA) and neuraminidase (NA), that have critical opposing functions in the IBV life cycle. The main function of HA is attachment and entry into host cells [5, 6], while NA facilitates release of new virions by cleaving sialic acids of cell surface receptors [7]. In addition, NA also enhances virus spread by enabling virus passage through mucus layers and prevents virion aggregation [8]. In turn, exploiting NA inhibition (NI) underpins anti-viral treatment available for influenza, with drug inhibition of NA by Oseltamivir (Tamiflu), Zanamivir (Relenza), or Peramivir (Rapivab) [9–12]. In addition, NA-specific antibodies have been associated with protection against influenza disease and viral replication [13–17]. As a result, NA is a promising target for the development of next-generation influenza vaccines against IAV and IBV [18]. The development of NA-based vaccines will require both an increased understanding of the antigenic evolution of NA, and accurate measurement of NI activity in human and animal serum. While considerable effort has been put into optimising and standardising NAI assays for IAV, little progress has been made for IBV.

Two commonly used assays to measure influenza virus NA and NI activity are the enzyme-linked lectin assay (ELLA) and the NA-Star/MUNANA assay. In the ELLA, the influenza virus NA cleaves sialic acids on fetuin, exposing the terminal galactose moiety, which can subsequently be detected using a lectin from *Arachis Hypogaea* (PNA), usually conjugated to Horseradish peroxidase (HRP) [19, 20]. The ELLA can detect NI by antibodies binding within the active site as well as those that bind proximally to the active site and block access to the NA substrate by steric hindrance [20]. However, it is well established that anti-HA antibodies can result in inhibition of NA activity via steric hindrance, due to the proximity of the NA and HA on the virion surface [21–23]. Consequently, the ELLA suffers from interference by anti-HA antibodies and can detect NI that is independent of NA-specific antibodies. The second NA assay, (MUNANA or NA-Star) utilises an acetylneuraminic acid (sialic acid) substrate linked with either fluorescent molecule (MUNANA) or chemiluminescent molecule (NA-Star) [24, 25]. Cleavage by the influenza virus NA releases the fluorescent or chemiluminescent reporter [24, 25]. Due to the small size of the MUNANA substrate, which allows it to easily access the enzymatic region on NA, HA interference by steric hindrance does not occur in the MUNANA assay [22]. However, for the same reason, the MUNANA assay only measures inhibition caused by antibodies that bind directly into the enzymatic active site of NA, and not those that might block access of NA to its substrate by steric hindrance [22]. As a result, the ELLA assay may capture a greater spectrum of NA-specific antibodies that can inhibit NA activity. However, HA interference must be overcome.

A variety of methods have been used to overcome interference by anti-HA antibodies for IAV and could potentially be applied to IBV. Firstly, recombinant NA (rNA) proteins can be used instead of virus, however, these may not be readily applicable to labs lacking expertise in recombinant protein expression or in experiments involving a large panel of NA antigens, which would need to be cloned, expressed and validated [26]. Secondly, recombinant influenza viruses that carry an non-human HA (e.g. H6 or H9) to which humans should have little to no antibodies have been commonly used in NA IAV serology studies [23, 26]. Reverse genetics have been used to generate recombinant IAV on the A/PR/8/34 backbone with a H6 [23] or H9 [26] HA and a chimeric NA consisting of IBV NA ectodomain linked to transmembrane domain and C-terminus from IAV. However, this approach could also prove impractical and laborious when a diverse panel of NA antigens is needed. Alternatively, splitting of influenza viruses could avoid the physical proximity of HA and NA, which is the main factor leading to NI mediated by anti-HA antibodies [21]. Another potential strategy is removal of anti-HA antibodies from sera samples, if appropriate HA antigens are available for absorption.

To understand and overcome HA interference for IBV, we assessed the extent of interference caused by anti-HA antibodies against IBV NA activity and examine several strategies to minimise it. We demonstrate that separation of HA and NA proteins on virions using Triton-X 100 detergent significantly reduces anti-HA antibody interference of IBV NA activity and characterise a panel of recombination IBV NA proteins (rBNAs). Using this optimised ELLA assay, we demonstrate discordant antigenic evolution of IBV NA and HA proteins over the last 80 years. These findings will aid serological surveys of IBV NA immunity as well as antigenic characterisation of the IBV NA on a larger scale.

## Results

### Treatment of serum for NI assays

We first assessed the use of Receptor Destroy Enzyme (RDE) to remove non-specific inhibition of the IBV NA as previously suggested for IAV [27]. As RDE is itself a neuraminidase, we confirmed that heat-inactivation for a minimum of 30 minutes was sufficient to completely inactivate all enzymatic activity of RDE in the ELLA assay **(Supplementary figure 1A)**. RDE treatment followed by 1 hour of heat inactivation did not impact the levels of anti-NA titres in human serum as determined by ELISA, but decreased inhibition observed in ELLA **(Supplementary figure 1B)**. Similarly, ELLA inhibition titres were lower in RDE and heat-inactivated serum from mice or ferrets compared to non-treated or only heat-inactivated samples. We therefore proceeded with overnight RDE treatment followed by 1 hr of heat-inactivation for subsequent ELLA experiments.

### Anti-HA antibodies mediate interference against NA activity of IBV

Although the interference of anti-HA antibodies against IAV NA activity has been well demonstrated [21, 28], less is known about the extent to which HA antibodies interfere with NA activity for IBV. ELLA was used to measure the NI activity of previously described anti-IBV- HA monoclonal antibodies (mAbs) CR8071 and CR8033 [29] **(Figure 1A)**, with two anti-IBV- NA mAbs 1G05 and 2E01 [30] used as positive controls. Both anti-HA mAbs showed NI activity in ELLA, although this varied across a panel of IBV isolates. While no NI was observed for either anti-HA mAbs against B/Florida/04/2006 or B/Brisbane/60/2008, both anti-HA mAbs showed an NI IC_50_>1ug/ml against B/Phuket/3073/2013 and B/Lee/1940. While some NI activity was observed against B/Malaysia/2506/2004 this remained less than 60%. NI activity was also observed within anti-B/Lee/1940 HA-specific serum with an NI IC_50_ titre against B/Lee/1940 virus of 1/602 dilution **(Figure 1B)**. Although the NI activity of NA-specific mAbs or serum was considerably higher than that of HA-specific mAbs/serum, these data clearly demonstrate that anti-HA antibodies can interfere with the enzymatic activity of IBV NA.

**Figure 1.**
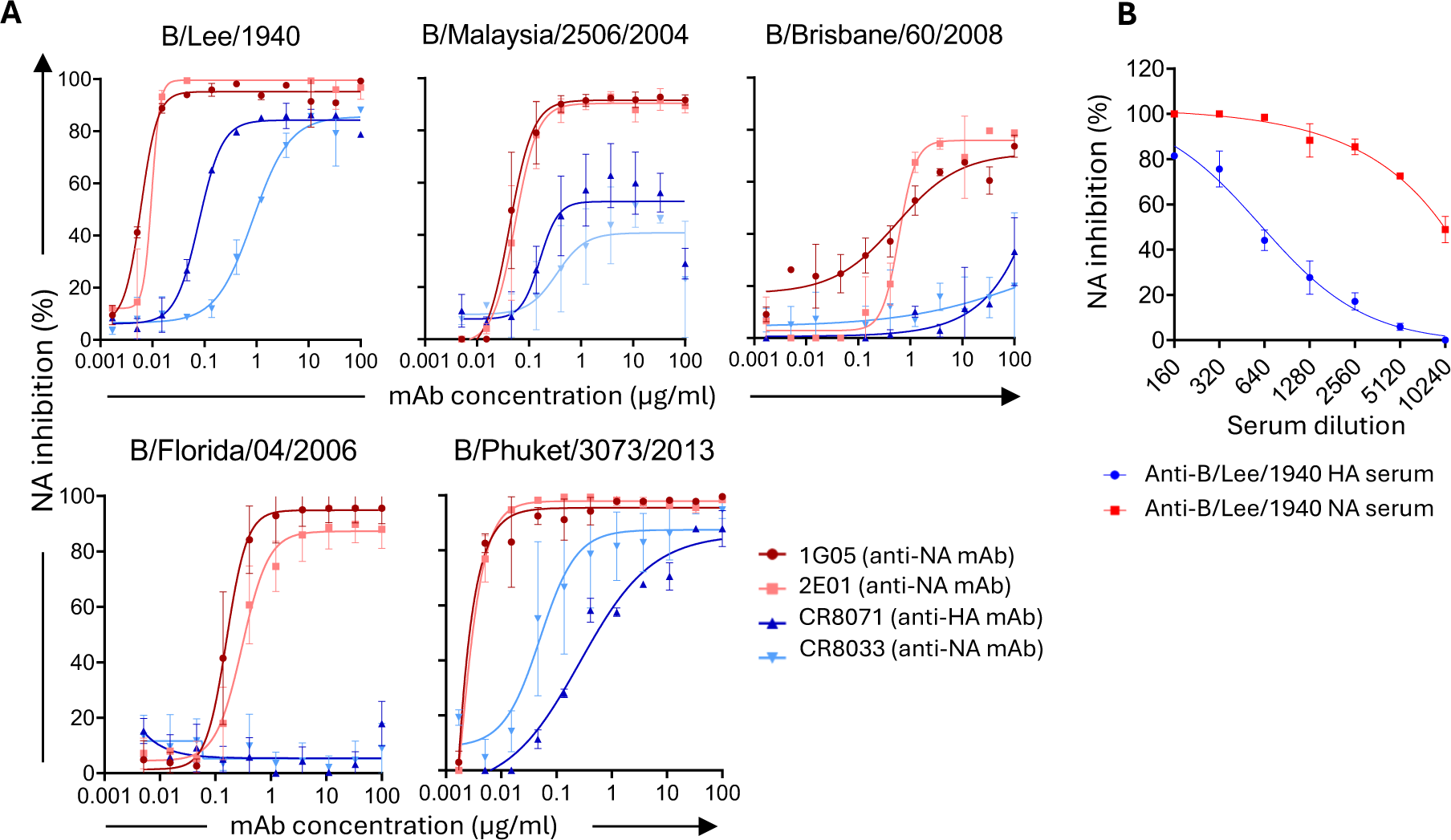
Anti-HA antibody-mediated interference against NA activity of IBV. (A) The NAI activity of two anti-HA mAbs (CR8071 and CR8033) was measured against 5 different influenza B viruses in an ELLA. Two anti-NA mAbs (1G05 and 2E01) were used as positive controls. **(B)** The NAI activity of anti-B/Lee/1940-HA serum was measured against B/Lee/1940 virus in an ELLA. Anti-B/Lee/1940-NA serum was used as a positive control. Virus input was standardized by EC90 value of virus stock NA activity. Throughout the figure the mean and standard deviation from 2 independent replicates is shown.

### Recombinant IBV NA proteins are enzymatically active and avoid HA interference

Recombinant IBV NA (rBNA) proteins provide a pathway to avoid NI interference caused by anti-HA antibodies and also side-steps propagation of viruses such as B/Yamagata lineages, where concerns exist about accidental reintroduction. We generated a panel of 7 rBNA proteins from the Ancestral, B/Victoria and B/Yamagata lineages, based on the stabilised design previously described by Ellis et al [31] **(Figure 2A)**. The rBNAs consist of the globular head domain of IBV NA linked the human Vasodilator-stimulated phosphoprotein (hVASP) tetramerization domain, a hexa-Histidine affinity tag (HisTag) and an AviTag for biotinylation. Soluble rBNA proteins were expressed and purified using polyhistidine-tag affinity chromatography and size-exclusion chromatography (SEC), with an expected size of approximately 50kD in SDS-PAGE (**Figure 2B)**. The conformational integrity of rBNAs was assessed by based upon binding to anti-NA mAbs (1G05 and 2E01) **(Figure 2C**) and enzymatic activity in ELLA and MUNANA **(Figure 2D)**. While these rBNA proteins may be useful for serological assessment of NI, they may not be practical for assays involving many different antigens which are required for antigenic cartography studies. We therefore assessed additional strategies to overcome HA interference.

**Figure 2.**
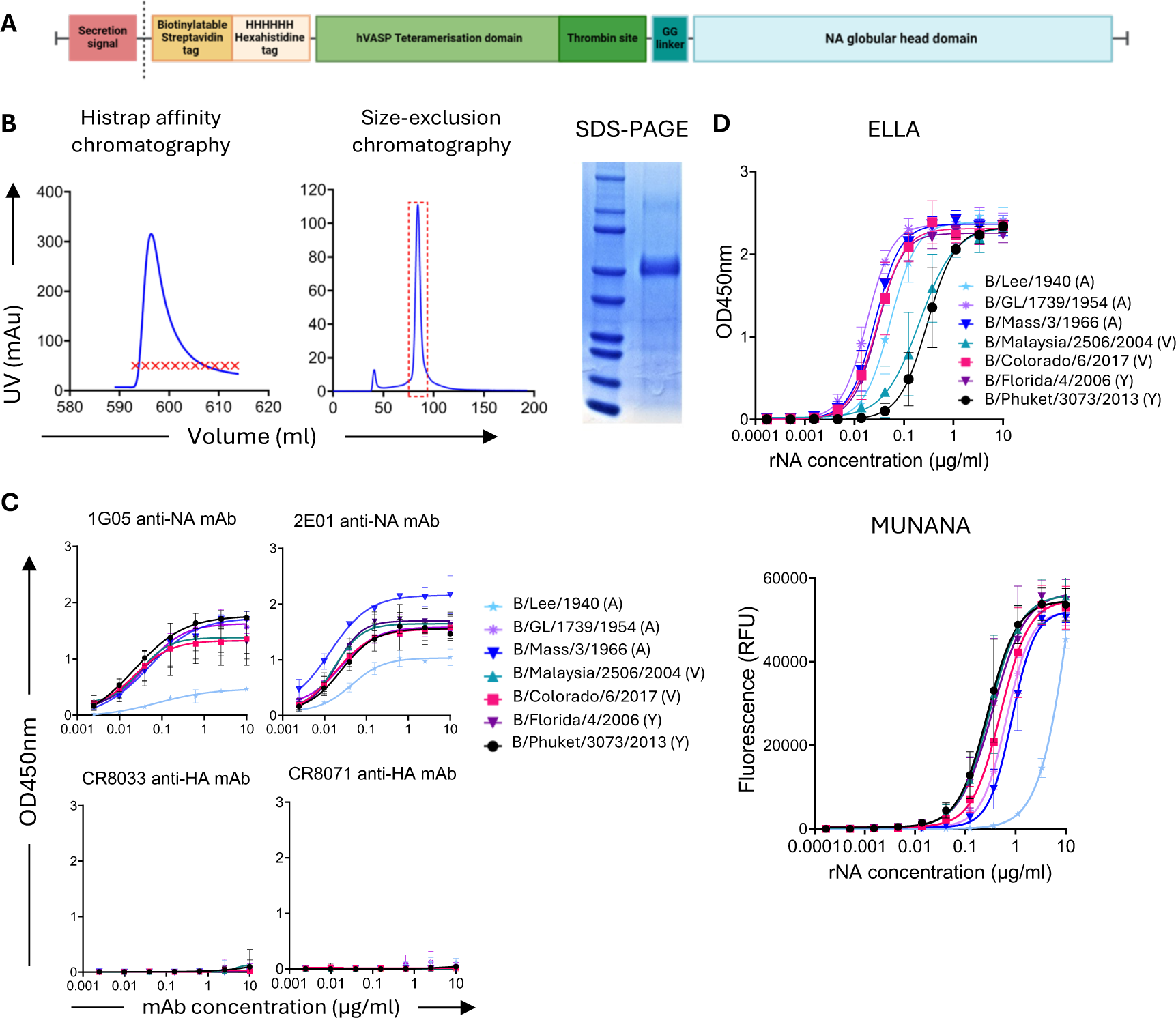
Expression and characterisation of recombinant IBV NA proteins. (A) Design of recombinant rBNA constructs. **(B)** rBNA was expressed by CHO cells and purified via 2 steps including Histrap affinity chromatography and size-exclusion chromatography, and then confirmed to have expected size by SDS-PAGE. **(C)** The conformation of rBNA was assessed by binding of known anti-NA mAbs (1G05 and 2E01) using ELISA. Two anti-HA mAbs (CR8071 and CR8033) were used as negative controls. **(D)** The enzymatic activity of rBNA was assessed by ELLA and MUNANA. For (C-D), the mean and standard deviation are shown (n=6, tested across 3 independent experiments).

### Incomplete depletion of anti-HA antibodies in animal and human sera

We next explored different strategies to avoid HA interference in assays using virus. Removing anti-HA antibodies from serum samples prior to ELLA could be a practical solution to limit HA interference. We initially attempted to deplete anti-HA antibodies by incubating serum samples with recombinant HA-coated plates prior to ELLA. We specifically used an anti-HA serum to directly determine the NI activity caused by anti-HA antibodies. The IC_50_ of the anti-HA serum decreased in a HA coating dose-dependent manner compared to sera in non-coated well **(Supplementary figure 2A)**. However, even at a high coating concentration of HA (40ug/ml), considerable residual HA interference of NA inhibition was observed.

We next generated cell lines stably expressing the HA from either B/Phuket/3073/2013 (B/Yamagata lineage) or B/Colorado/06/2017 (B/Victoria lineage) **(Supplementary figure 2B)**. Firstly, we assessed the depletion of anti-HA antibodies in mouse anti-sera (anti-B/Phuket/3073/2013 and anti-B/Colorado/06/2017) **(Supplementary figure 2C)** or human sera (on day 7 post-vaccination) **(Supplementary figure 2D)**. Incubation of sera with increasing number of HA-expressing cells resulted in reduction of homologous anti-HA antibodies, measured by ELISA, relative to non-treated samples **(Supplementary figure 2C- D)**. However, we also observed some non-specific depletion of HA antibodies in samples treated with WT 293T (HA null) cells as well as in the levels of NA-specific antibodies. We further examined whether this strategy would efficiently reduce anti-HA antibodies heterologous to the depleting HA strain. To that end, human sera samples were treated with either B/Colorado/06/2017-HA 293T cells or B/Phuket/3073/2013-HA 293T cells or a mix of both and then residual antibodies against different IBV HA were assessed by ELISA. Although there was a reduction of HA antibodies against B/Florida/04/2006 and B/Mass/3/1966, depletion was less effective against B/GL/1739/1954 and B/Allen/1945 **(Supplementary figure 2E).** Similarly, depletion of heterologous anti-HA antibodies from different mouse anti- sera was limited **(Supplementary figure 2F)**. In conclusion, given the limited effects on depleting heterologous HA antibodies and the non-specific impact on NA antibodies, depletion of anti-HA antibodies from either human sera or animal antisera may be of limited utility in overcoming HA interference.

### Triton-X 100 efficiently reduces HA interference by separating HA and NA from virion

Given that anti-HA antibodies interfere with NA activity by steric hindrance [21, 22], another strategy to avoid the NI caused by anti-HA antibodies is by separating the HA interference arises from co-localisation of HA and NA molecules on the virion surface [22]. We therefore assessed separation of IBV HA and NA from virions using detergent (Tween-20 or Triton-X 100) or ether disruption. A capture ELISA using an anti-HA stem antibody (CR9114) to capture HA and anti-HA or anti-NA antibodies for detection was used to determine the association of HA and NA in the presence of Tween-20, Triton-X 100, or using Ether-split IBV **(Figure 3)**. Two IBV isolates (B/Lee/1940 and B/GL/1739/1954) were tested. Both Tween-20 or ether-treated viruses failed to dissociate HA and NA proteins. In contrast, when using Triton-X 100 we could not detect any anti-NA signal after capturing HA antigens, suggesting efficient dissociation of HA and NA. This result suggests that Triton-X 100 separates NA and HA proteins from influenza B virions.

**Figure 3.**
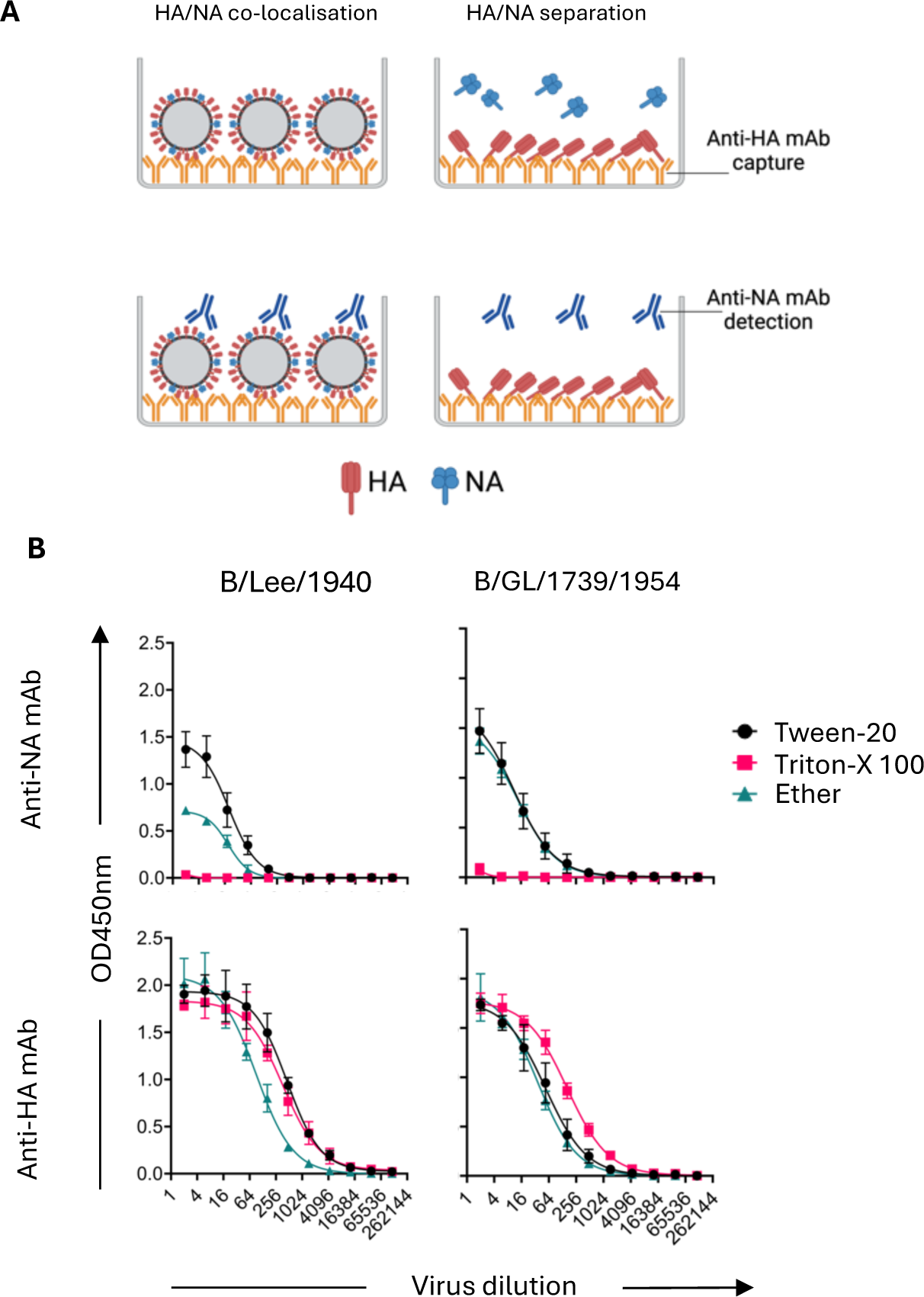
Separation of HA and NA proteins from virion by Triton-X **100**. B/Lee/1940 and B/GL/1739/1954 viruses were treated with either 0.5% Tween- 20 or 0.5% Triton-X 100 or Ether. The amount of NA and HA proteins was determined by capture ELISA, starting from 1:2 dilution of virus. The mean and standard deviation are shown (n=4, tested across 2 independent experiments).

To determine if dissociation of HA and NA by Triton-X-100 was sufficient to overcome HA interference, we compared the use of ether-split IBV or the addition of 0.5% Triton-X to the standard ELLA protocol (0.5% Tween-20). These conditions were tested using 3 IBV strains (B/Lee/1940, B/GL/1739/1954, and B/Phuket/3073/2013) in combination with anti-HA or anti-NA mAbs **(Figure 4A)**. NI activity of anti-HA mAbs against B/Lee/40 was increased when using ether-split virus compared to Tween-20 only. In contrast, NI activity of anti-HA mAbs was only evident at the highest concentration of mAbs (100ug/ml) and at very low levels (∼20%) when using Triton-X. Similarly, although HA interference was generally lower against B/GL/1739/1954, it was reduced in the presence of Triton-X 100. Consistently, HA interference against B/Phuket/3703/2013 was considerably reduced by either the use of ether-treated IBV or the addition of Triton-X 100. We further assessed the effect of Triton-X by using mouse anti- sera specific for IBV HA antigens (B/Lee/1940-HA, B/GL/1739/1954-HA, and B/Phuket/3073/2013-HA) **(Figure 4B).** While NI activity was observed for these anti-HA specific sera in the standard buffer (Tween-20) and was increased when using ether-treated B/Lee/1940-HA and B/GL/1739/1954, NI activity of HA anti-sera was markedly reduced when Triton-X 100 was added. These data suggest that addition of Triton-X 100 efficiently overcomes interference by anti-HA antibodies with little impact on anti-NA antibodies.

**Figure 4.**
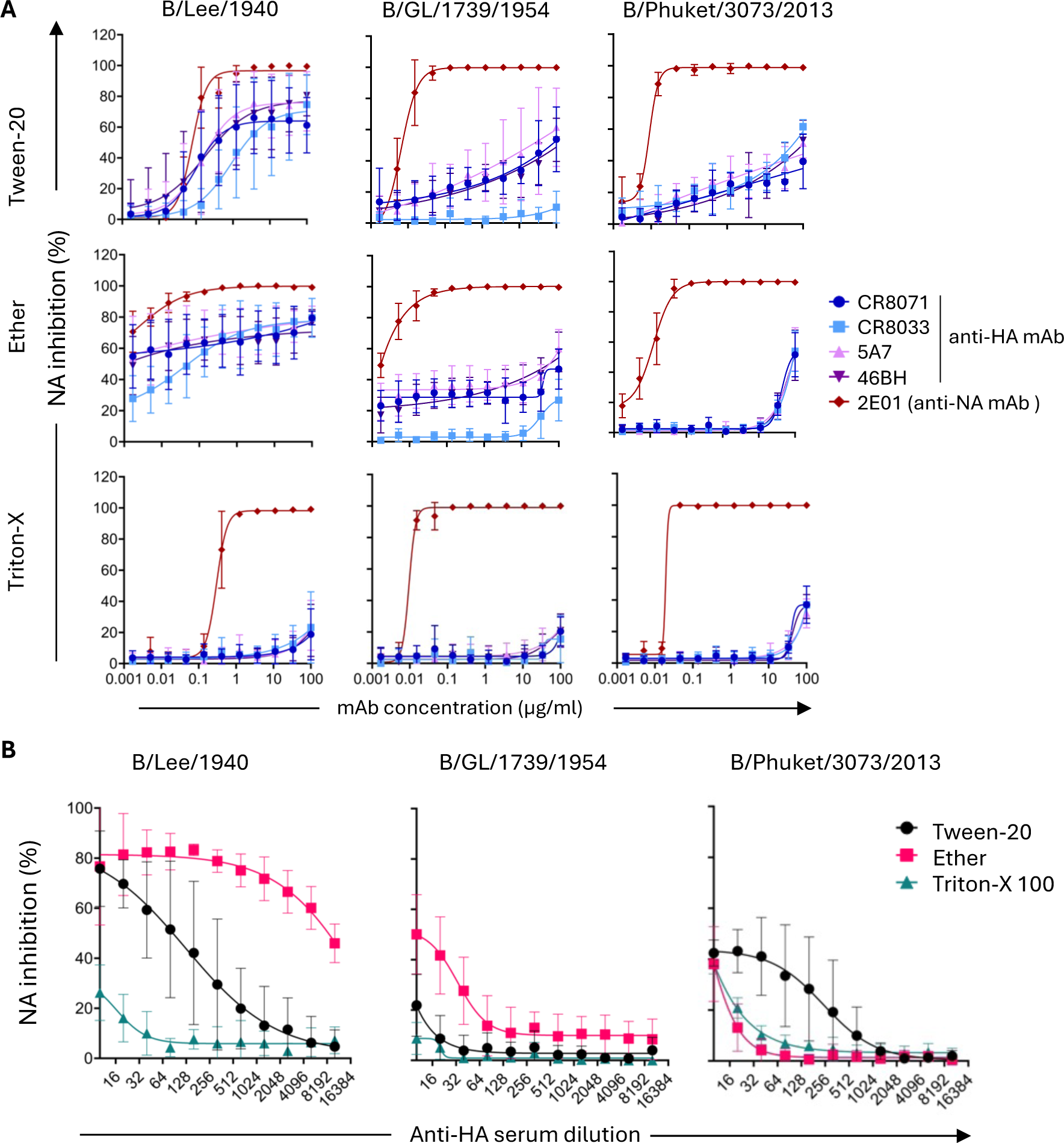
Triton-X efficiently reduces NA inhibition mediated by anti-HA antibodies **(A)** 0.5% Tween-20 or 0.5% Triton were added to the assay buffer or IBV were ether-split before testing in an ELLA using four different anti-HA mAbs (CR8071, CR8033, 5A7, and 46B8). Two anti-NA mAbs (1G05 and 2E01) were used as positive controls. **(B)** NAI activities of mouse anti-HA sera against IBV isolates were determined in an ELLA with either 0.5% Tween-20, 0.5% Triton, or ether-treated IBV. For (A-B), virus input was standardized by EC90 value of virus stock NA activity. The mean and standard deviation are shown (n=5, tested across 3 independent experiments).

Finally, we investigated the effects of Triton-X-treated viruses on HA interference in human sera. To that end, we treated human sera (obtained 7 days after seasonal influenza vaccination) with 293T cells transiently expressing the NA from B/Florida/04/2006 or WT cells, resulting in depletion of NA-specific antibodies **(Figure 5A)** but not HA-specific antibodies, measured by ELISA **(Figure 5B)**. We next tested the NA-293T-treated (NA ab^-^/HA ab^+^), WT- 293T-treated (NA ab^+^/HA ab^+^) and untreated samples (NA ab^+^/HA ab^+^) in an ELLA with Tween- 20 or with Triton-X-100 against B/Phuket/3073/2013 and B/Lee/1940. Serum samples were tested starting at a dilution of 1/800 at which there were high level of anti-HA antibodies **(Figure 5B)** and little residual anti-NA antibodies left **(Figure 5A)**. In the standard ELLA buffer (Tween-20) low levels of residual NI could be observed for both viruses after depletion of NA-specific antibodies. However, in the presence of Triton-X, NI in NA-antibody-depleted sera was completely removed. While the residual NI activity observed in NA-antibody-depleted sera and Tween-20 treated virus was on average low (≤20%), it ranged from 8-34% across the 8 individuals tested. In contrast it was consistently below 10% across all donors in the presence of Triton-X **(Figure 5C)**. Overall, inclusion of Triton-X 100 in the ELLA assay buffer overcomes interference mediated by anti-HA antibodies, as showing in the context of anti-HA mAbs, mouse anti-sera, and human sera.

**Figure 5.**
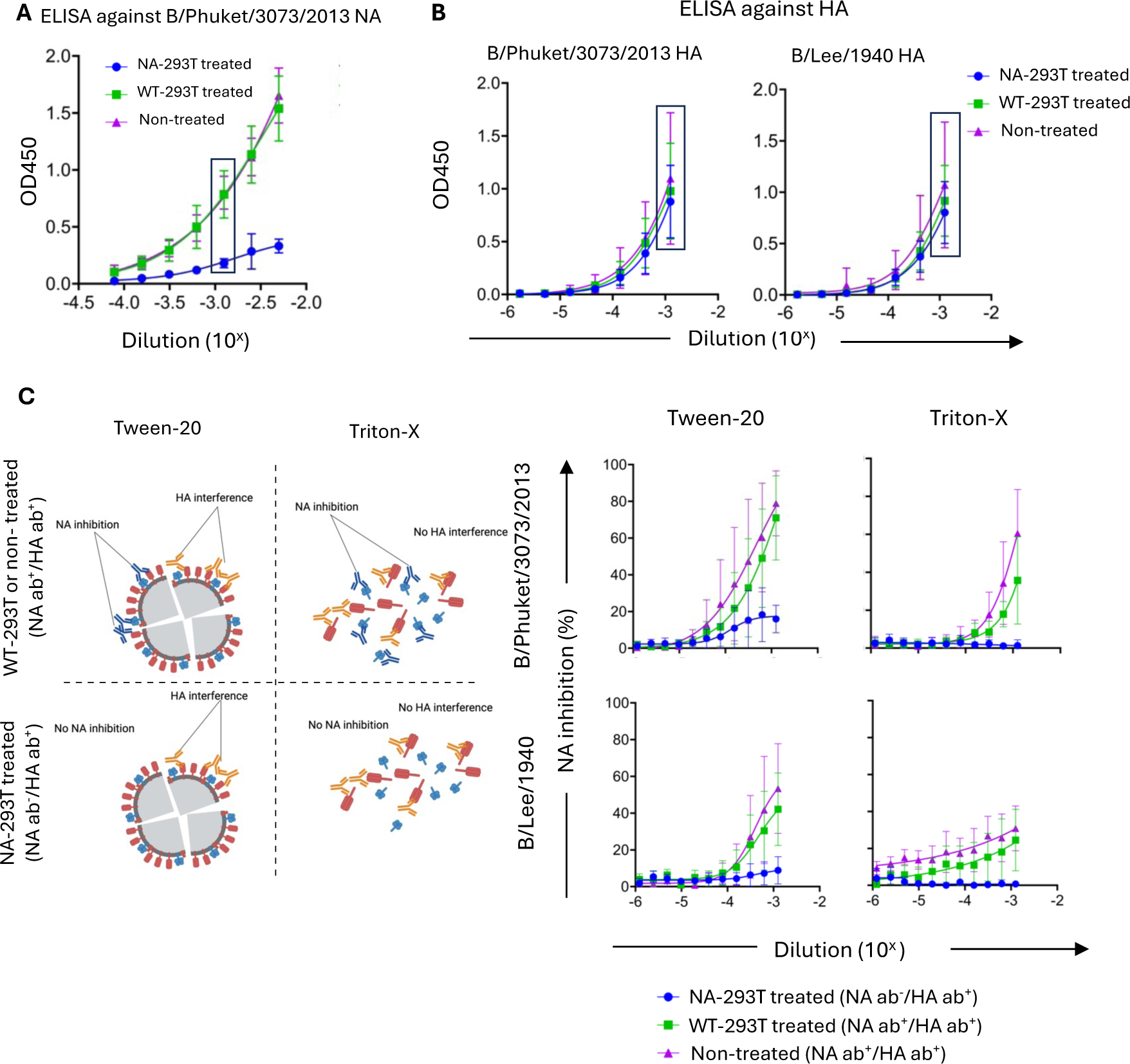
Triton-X efficiently removes interference mediated by anti-HA antibodies in human sera. (A) Human sera were incubated with 4x10^6^ of B/Florida/04/2006 NA-expressing 293T cells. Sera samples were incubated with 4x10^6^ of WT 293T cells or left non-treated as negative controls. Depletion of anti-NA antibodies was determined by ELISA against B/Phuket/3073/2013 NA (starting from 1/200 dilution of sera). **(B)** The amount of anti-HA antibodies after depletion was determined by ELISA against B/Phuket/3073/2013 and B/Lee/1940 HA (starting from 1/800 dilution of sera). **(C)** NAI activities of anti-NA antibodies depleted human sera against IBV isolates were determined in an ELLA with either 0.5% Tween-20 or 0.5% Triton IBV. Virus input was standardized by EC90 value of virus stock NA activity. For (A-C), the mean and standard deviation are shown (n=8 donors, tested across 2 independent experiments).

### NA inhibition titres of sera against Triton-X-treated viruses and recombinant rBNA proteins are positively correlated

To confirm the observations made using Triton-X-treated viruses, we next correlated NI titres of ferret antisera measured in ELLA using matched rBNA proteins and Triton-X-treated viruses. Seven different ferret antisera were assessed against a panel of 7 homologous rBNA proteins and Triton-X-treated viruses including B/Lee/1940 (Ancestral lineage - A, with regards to HA), B/GL/1739/1954 (A), B/Mass/3/1966 (A), B/Malaysia/2504/2004 (Victoria lineage – V), B/Colorado/06/2017 (V), B/Florida/04/2006 (B/Yamagata – Y) and B/Phuket/3073/2013 (Y) **(Figure 6A)**. Using either rBNA or Triton-X-treated viruses we observed that anti-sera raised against Ancestral viruses primarily showed cross-reactivity towards Ancestral but not B/Vic or B/Yam isolates, and vice versa. The only inconsistency between the two was that the Triton-X-treated B/Phuket virus was cross-recognised by Ancestral antisera while the B/Phu rBNA was not. Overall, however, the NI titres (IC_50_) determined using matched rBNA proteins and Triton-X-treated viruses were positively correlated (r=0.81, p<0.0001) **(Figure 6B)**.

**Figure 6:**
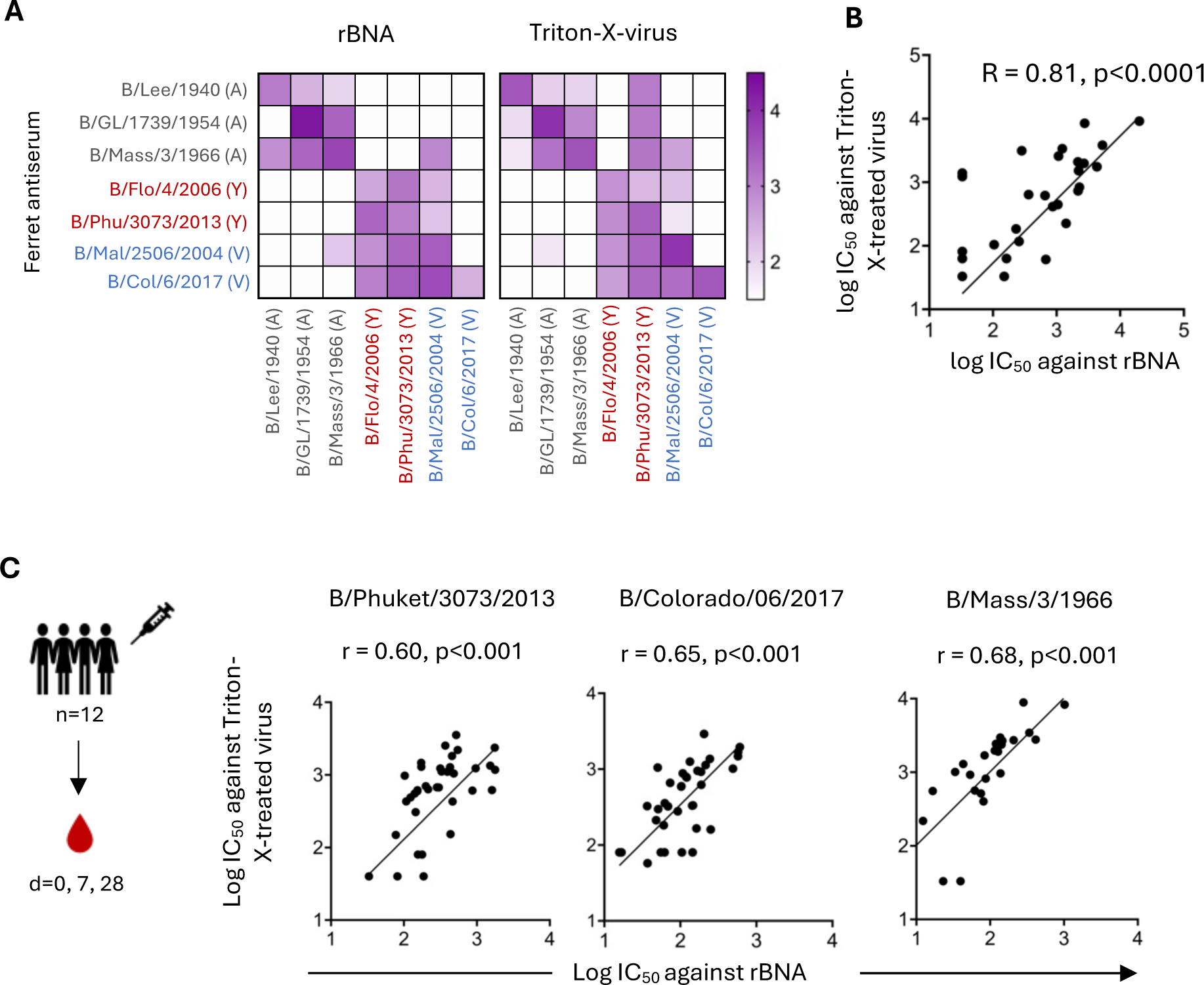
Positive correlation between NA inhibition titres of sera against Triton-X- treated viruses and recombinant IBV NA proteins. (A-B) The NI activities of 7 different ferret antisera against homologous and heterologous rBNAs and Triton-X 100-treated viruses were tested. The IC_50_ values were obtained and mapped (A). The Spearman correlation between all titrations was assessed (n=49) (B). **(C)** The NI activities of human sera were tested using Triton-X 100-treated IBV and rNA proteins for three strains. The sera were collected from 12 healthy donors at baseline and 7 days and 28 days post- vaccination. The Spearman correlation between all titrations for each virus was assessed (n=36, from 12 donors). The virus input was standardized at EC_90_ value for ELLA with ferret antisera and EC_70_ for ELLA with human sera. The rNA protein input was standardized at EC_50_ value for both test with ferret antisera and human sera.

We next assessed NI against B/Phuket/3073/2013, B/Colorado/06/2017, and B/Mass/3/1966 in 36 human sera samples using matched rBNA and Triton-X-treated viruses. The samples were obtained from 12 donors before and 7 or 28 days post-vaccination with the 2022 quadrivalent influenza vaccine (containing B/Phuket/3073/2013 and B/Colorado/06/2017). Our aim was not to determine NA responses to vaccination (which will vary depending on NA content), but rather to assess NI in the context of varying levels of anti-NA and anti-HA antibodies. Consistent with the positive correlation seen in ferret anti-sera, we observed positive correlations between NI titres of human sera using rBNA and Triton-X- treated viruses (B/Phuket/3073/2013 r=0.60, p<0.001; B/Colorado/06/2017 r=0.65, p<0.001; B/Mass/3/1966 r=0.68; p<0.001) **(Figure 6C)**. These data further demonstrate that using Triton-X to dissociate the HA and NA is an effective approach to determine NI without anti-HA antibody-mediated interference.

### Discordant evolution of the IBV HA and NA between 1940-2017

While the antigenic diversification of the IBV HA into two antigenically distinct lineages has been well-established [32, 33], the antigenic diversity of the IBV NA is less understood. To address this, we utilised the Triton-X-based ELLA to probe the antigenic evolution of the IBV NA. Consistent with previous findings [32, 33], phylogenetic analysis of the IBV HA from 13 isolates spanning 1940-2017 clearly indicated the divergence of the B/Victoria and the B/Yamagata lineages from the Ancestral lineage (**Figure 7A**). This was not evident in the NA phylogeny using the same isolates (**Figure 7B**). Rather, B/Vic/2/1987 is ancestral to the clade containing all other B/Vic and B/Yam sequences and the B/Vic/2/1987 and B/Yam/16/1988 are not ancestral to the B/Vic and B/Yam lineages, respectively, as seen in the HA. In addition, B/Sichuan/379/1999 (belonging to B/Yam based on HA) was ancestral to B/Malaysia/2506/2004 and subsequent NAs from B/Victoria-like isolates, consistent with the reassortment of the NA segment between the two lineages in the early 2000s [33–35].

**Figure 7.**
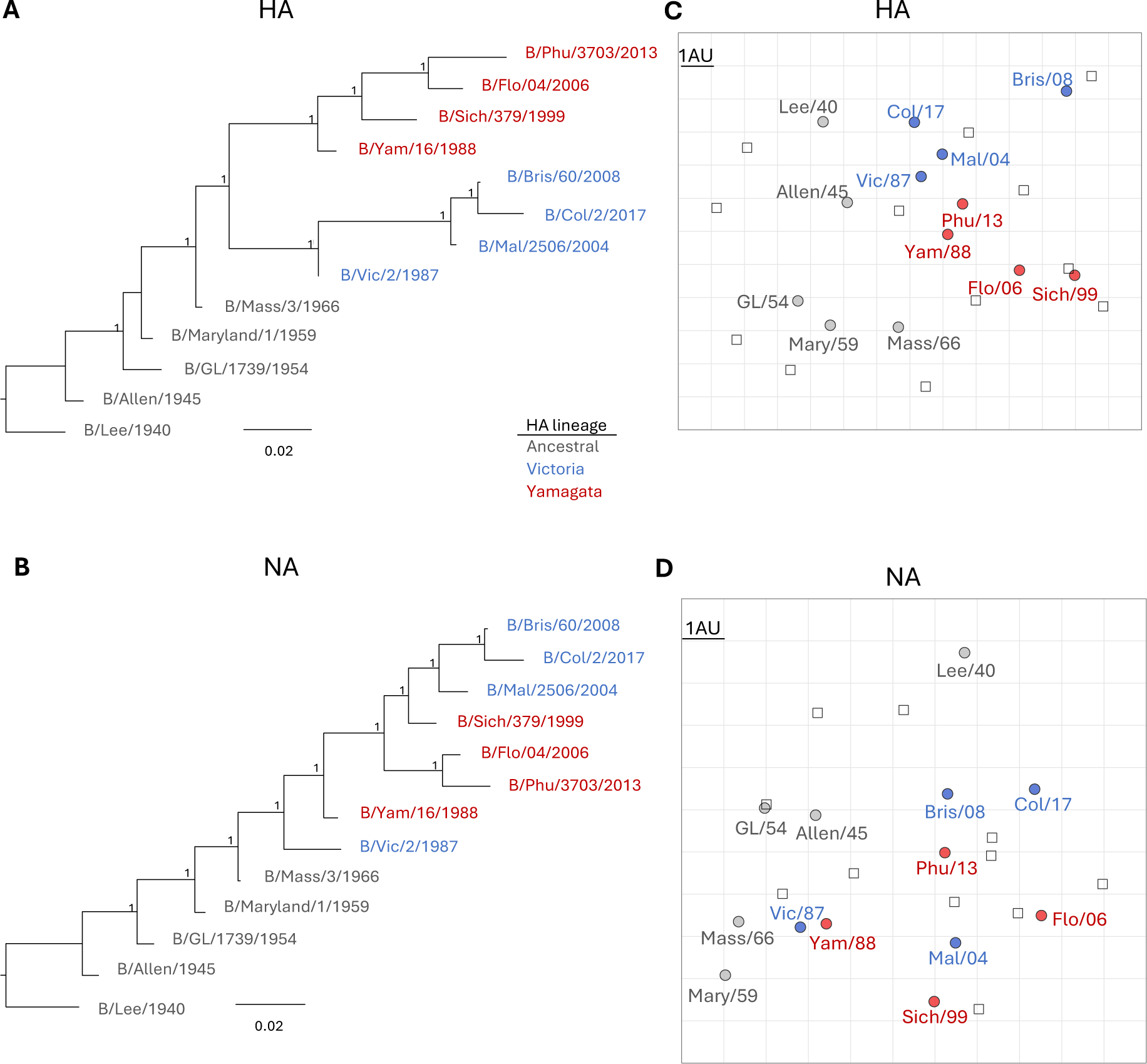
**Antigenic characterisation of the IBV HA and NA**. **(A-B)** Phylogenetic tree of 13 IBV isolates from 1940-2017 for the HA (A) and NA (B)**. (C-D)** Phylogenetic tree of 13 IBV isolates from 1940-2017 for the HA (C) and NA (D). The circles represent position of the isolates, and the squares represent positions of the ant-sera. Each square represents 1 antigenic unit (AU), equivalent to a 2-fold difference in titres. Isolates are color-coded based on the HA lineage throughout the figure. **(E)** Correlation between pairwise antigenic distance of the HA and NA between consecutive isolates (eg between B/Lee/1940 and B/Allen/1954, between B/Allen/1945 and B/GL/1954).

Antigenic cartography of the HA from these 13 isolates using ferret anti-sera in HAI assays (**Figure 7C**), demonstrated separation of Ancestral isolates from the 1940s (B/Lee/1940 and B/Allen/45) and those from the 1950s and 1960s (B/GL/1954, B/Maryland/1/1959 and B/Mass/2/66) (average pairwise distance of 4.6 antigenic units – AU). The Ancestral viruses from the 1950s and 1960s were also antigenically separate from isolates of either the B/Victoria or B/Yamagata lineages (average pairwise distance of 6.1 AU). B/Victoria and B/Yamagata isolates also demonstrated antigenic separation as expected (average pairwise distance of 4.2 AU). In addition, we observed antigenic change within each lineage. For example, B/Malaysia/2506/2004 was 4.2 AU away from B/Brisbane/60/2008 which in turn was 4.6 AU away from B/Colorado/2/2017. Similarly, B/Yamagata/16/1988 was 4 and 2.5 AU away from B/Sichuan/379/1999 and B/Florida/04/2006 respectively, and these were in turn 2 and 2.6 AU away from B/Phuket/3703/2013. These antigenic data are overall consistent with previous antigenic characterisation of the IBV HA [33, 36] and the periodic updates of vaccine strain selection for each IBV lineage.

We next applied antigenic cartography to the IC_50_ NI titres from a Triton-X-based ELLA using the same antigens and anti-sera (**Figure 7D**). The NA antigenic map did not reveal the same antigenic separation as for the HA. For example, B/Allen/45 and B/GL/1954, which were antigenically separate on the HA map (3.3 AU) were antigenically close on the NA map (1.2 AU). Similarly, while B/GL/1954 was antigenically similar to B/Maryland/1/1959 on the HA map (1.2 AU) it was antigenically separate on the NA map (4.1 AU). In addition, while B/Victoria/2/1987 and B/Yamagata/16/1988 were antigenically separate from B/Mass/2/66 on the HA map (4.6 and 3.2 AU respectively) they were antigenically close on the NA map (1.5 and 2.1 AU respectively). Consistently, the B/Victoria/2/1987 and B/Yamagata/16/1988 isolates were 1.9 AU on the HA map but only 0.6 AU away in the NA map. NA from subsequent co-circulating isolates from each 2 lineages were also antigenically close to each other (eg. B/Sichuan/379/1999 and B/Malaysia/2506/2004; or B/Phuket/3703/2013 and B/Colorado/2/2017). Overall, our paired antigenic characterisation of the IBV HA and NA demonstrate the discordant evolution of these surface glycoproteins.

## Discussion

Interference by anti-HA antibodies in ELLA has been well established for IAV and various strategies have been explored to overcome it. This includes the generation of H6Nx [23] or H9Nx [26] viruses. Although this method can reduce the effect of anti-HA antibodies, it might be laborious and time-consuming if a variety of different NAs need to be assessed. There may also be HA stem antibodies cross-reactive between human H1 and animal H6 IAV that may still contribute to low levels of HA interference [23]. Alternatively, the addition of Triton-X-100 can overcome HA interference caused by antibodies against the IAV HA [22]. HA interference in the context of IBV has not been similarly investigated. Here we demonstrate that addition of Triton-X-100 is sufficient to overcome HA interference for IBV and apply this to probe the antigenic evolution of the BNA.

In our experiments, the addition of Triton-X-100 was sufficient to separate the IBV HA and NA and almost completely overcame HA interference caused by anti-HA monoclonal antibodies, or human and animal serum. We speculate that the effects of Triton-X 100 are due to its ability to solubilize membranes and extract proteins [37, 38], although it is unclear why this was not achieved by Tween-20. Consistently, we could not co-detect HA and NA in the presence of Triton-X-100, while HA and NA could still be co-detected in ether-treated viruses, which in some cases demonstrated increased HA interference. Ether-treatment increases the sensitivity of HAI against IBV, presumably by exposing HA domains that are otherwise occluded in whole virions. This may result in increased binding of HA antibodies thus explaining the increased HA interreference observed. The addition of Triton-X-100 to overcome HA interference for IBV is consistent with previous studies in IAV [22]. Thus, the addition of Triton-X 100 is a simple modification to the standard ELLA protocol that can be readily applied and standardised across laboratories and does not require any particular expertise or reagents like recombinant protein expression or the generation of chimeric IAV/IBV.

We examined additional strategies to avoid HA interference for IBV. The efficiency of depleting of anti-HA antibodies by HA-expressing 293T cells was strain-specific and not applied to all tested strains. We also noted non-specific depletion of NA antibodies. Therefore, this approach may be of limited utility. We also expressed and validated a panel of 7 rBNA proteins that appear antigenically intact and are enzymatically active. While these may be useful as they completely avoid HA interference, rBNAs may be less useful for large-scale antigenic characterisation where viral isolates are more easily obtained or be suitable reagents in cases where viruses are unavailable or undesirable. In either case, ELLA with rBNA and Triton-X-treated virus were highly correlated in our analyses.

As there is limited experimental evidence for the extent of BNA antigenic evolution, we used the Triton-X-100 virus-based ELLA to probe the antigenic evolution of the IBV NA between 1940-2017. Although our antigenic cartography is limited to 13 isolates, we can clearly observe antigenic differences between IBV isolates of this time period. In addition, by comparing the patterns of antigenic evolution of IBV HA and NA glycoproteins, we observed that antigenic changes in one surface glycoprotein were discordant with antigenic change in the other surface glycoprotein, consistent with observations for H1N1 and H3N2 [39]. Further antigenic characterisation and sequence analyses may reveal BNA antigenic clusters and the mutations that underpin them and how they relate to HA antigenic clusters.

Despite finding evidence of antigenic drift of the IBV NA, limited diversification of the NA into antigenically distinct co-circulating lineages was observed in our antigenic cartography, unlike the B/Yamagata and B/Victoria lineages of the IBV HA. This is exemplified by the antigenic similarity of the NAs from the HA lineage-defining B/Victoria/2/1987 and B/Yamagata/16/1988 isolates. IBV NA diversification into distinct lineages may be generally limited by inter-lineage re-assortment of the NA segment. Indeed, in the early 2000s, viruses bearing the B/Victoria HA acquired a NA segment originating from the B/Yamagata lineage [34, 35]. This is evident in both the phylogeny of the NA (Figure 7B) [32, 34, 35] as well as the NA antigenic map, with the B/Malaysia/2506/2004 (V) NA being antigenically close to the NA from B/Sichuan/379/1999 (Y) and B/Florida/04/2006 (Y). As NA segments have been occasionally exchanged between the two lineages [32, 34–36], additional antigenic characterisation is needed to fully understand the antigenic dynamics of the BNA. In addition, since HA lineage-specific immunological imprinting has been observed [40] and is thought have shaped IBV epidemiology over the last 20 years [41], further antigenic characterisation of the BNA and antibody landscape analyses could also determine if NA imprinting occurs and contributes to birth cohort-specific IBV susceptibility.

A more detailed mapping of both the IBV HA and NA, and integration with epidemiological and sequence data is required to understand the antigenic dynamics of IBV and reveal the relative contribution of IBV HA and NA antigenic evolution to the spread of IBV. Such large-scale serological analyses can now be facilitated by the use of the Triton-X-100 virus-based ELLA without HA interference.

## Acknowledgements

We thank the participants for their generous involvement and provision of samples. We are grateful to the WHO Collaborating Centre for Reference and Research on Influenza in Melbourne for the provision of ferret anti-sera. The work has been generously supported by the Morningside Foundation and by Australian National Health and Medical Research Council Investigator grants (1195698 to M.K. and 1173433 to A.K.W.) This research was funded in whole or part by the National Health and Medical Research Council (1195698). For the purposes of open access, the author has applied a CC BY public copyright licence to any Author Accepted Manuscript version arising from this submission

## Author contributions

M.K. designed and supervised the study. T.H.T.D. performed the experiments. T.H.T.D., M.W. and M.K. analysed data. A.K.W provided samples critical for the study. T.H.T.D., A.K.W and M.K. contributed to drafting of the manuscript. All authors reviewed the final version of the manuscript.

## Competing interest

M.K. has acted as a consultant for Sanofi group of companies. The other authors declare no competing interests.

## Materials and methods

### Viruses & reagents

Viruses were either propagated in eggs (B/Lee/1940 (Ancestral – A), B/GL/1739/1954 (A), B/Mass/3/1966 (A), B/Florida/04/2006 (Yamagata – Y)) or MDCK cells (B/Malaysia/2506/2004 (Victoria – V), B/Brisbane/60/2008 (V), B/Colorado/06/2017 (V) B/Phuket/3073/2013 (Y)). Recombinant HA proteins and monoclonal antibodies were produced in house as previously described [42]. Recombinant NA (rNA) constructs are designed based on the study of Ellis et al [31]. The sequence of the NA was obtained from sequencing of viral stocks. The structure of rNA consists of the ectodomain (head domain only) of NA linked with a hexa-Histidine affinity tag (HisTag) and a biotinylatable AviTag (StrepTag). The soluble recombinant NA proteins are expressed in the form of tetramer by Chinese Hamster Ovary (CHO) cells and purified via 2 steps including polyhistidine-tag affinity chromatography and size-exclusion chromatography (SEC). The final protein product was assessed by SDS-PAGE to confirm the correct size.

### Human serum samples

Samples from adults were collected under study protocols that were approved by the University of Melbourne Human Research Ethics Committee (2056689). All participants provided written informed consent in accordance with the Declaration of Helsinki. Twelve healthy adults were vaccinated with the 2022 quadrivalent influenza vaccine containing following viral strains: A/Victoria/2570/2019(H1N1) pdm09-like, A/Darwin/9/2021(H3N2)-like, B/Austria/1359417/2021-like (B/Victoria lineage), B/Phuket/3073/2013-like (B/Yamagata lineage). Blood samples were collected at baseline (day 0), day 7, and day 28 post-immunisation. Sera were isolated and cryopreserved at -80◦C for future use.

### Animal anti-sera

Mouse antisera were generated by infection with live IBV or vaccination with recombinant HA/NA proteins. Female or male C57BL/6L mice aged 6–12 weeks were obtained from the Animal Resources Centre (Perth, Western Australia) or the Biological Research Facility in the Department of Microbiology and Immunology at the University of Melbourne (Melbourne, Victoria). All animal work was conducted in accordance with guidelines set by the University of Melbourne Animal Ethics Committee (ethics approval number 21799). For infection, mice were infected intranasally with 10^3^ plaque forming units (pfu) of IBV at the indicated doses diluted in 50μl of PBS under isoflurane anaesthesia. Blood was collected via the cardiac route (terminal) at 28-35 days post infection. For vaccination, mice were vaccinated intramuscularly with 5ug of protein in Addavax adjuvant (InvivoGen) (1:1 v/v) in a final volume of 50ul in each hind quadricep under light isoflurane anaesthesia. Animals were boosted 3 weeks after the priming and blood was collected at 14 days post-boost. Ferret anti-sera were provided by the WHO Collaborating Centre for Reference and Research on Influenza in Melbourne.

### Generation of stable HA-expressing cell lines and NA-expressing 293T cells

The B/Phuket-HA and B/Colorado-HA genes were coned into the pHAGE2 vector [43]. Next, the pHAGE2-HA plasmids were co-transfected with pRC-CMV-revb, pMD2-G (VSG-G), pHDM-Hgm2, and pHDM-tat1b into 293T cells. After 48 hours of transfection, the virus supernatant was harvested and frozen down. Transduction of lentiviruses into desired 293T cell line was conducted for 4-5 days. Transduced cells were then bulk sorted for top 5-10% of HA expression and regrown in fresh media. Expression of HA protein was confirmed by flow cytometry. For transient expression of NA, 293T cells were transfected with 1ug of a plasmid expressing the B/Florida/04/2006 NA under a CMV promoter using Lipofectamine^TM^ 3000 transfection reagent kit (Invitrogen). Cells were harvested and used for absorption after 48 hours of transfection.

### Serum treatment

For all experiments, sera were first treated with RDE (Denka Seiken) according to the manufacturer’s instructions, followed by heat-inactivation at 56°C for 1 hour. For absorption of anti-HA antibodies in sera by HA-expressing 293T cells, 50ul of RDE-treated sera samples were incubated with different number of HA-expressed 293T cells ranging from 0.5x10^6^ to 4x10^6^ cells in 96-well plate for 1 hour at room temperature with shaking. For absorption of anti-NA antibodies, 50ul of RDE-treated sera samples were incubated with to 4x10^6^ of NA- expressed 293T cells in 96-well plate for 1 hour at room temperature with shaking. Sera were treated with 4x10^6^ of WT 293T cells in 96-well plate for 1 hour at room temperature with shaking as negative controls.

### ELLA for NA activity

For all ELLAs, Maxisorp 96-well plates (Thermo Fisher) were coated with 100μl of Fetuin from fetal bovine serum (Sigma) at concentration of 25μg/ml, at 4◦C overnight. An assay buffer with 33 uM MES, 4 mM CaCl2, pH 6.5, 1% BSA, 0.5% detergent (either Tween- 20 or Triton-X) in distilled water was used. Horseradish peroxidase-conjugated arachis hypogaea lectin (PNA-HRP) (Eylabs) was diluted in PNA-diluent (33 uM MES, 4 mM CaCl_2_, pH 6.5, 1% BSA in water) at concentration of 1μg/ml. A sashing buffer PBS with 0.05% Tween- 20 (PBS-T) was used for all washing steps.

For the measurement of NA activity, viruses were serially diluted 2-fold in assay buffer (starting from 1:2 dilution). Alternatively, rNA proteins were serially diluted 3-fold in assay buffer (starting from 10μg/ml). Next, 25μl of assay buffer followed by 25μl of diluted virus or protein were added into Fetuin-coated Maxisorp plate and incubated at 37°C for 16-18 hours. Then, 100μl of PNA-HRP was added and incubated for 2 hours at room temperature. Finally, 100μl of TMB (Sigma) substrate was added for 3 minutes and the reaction was stopped by adding 50μl of stop solution (0.16M sulphuric acid). Between each step, plates were washed 6 times with PBS-T. Plates were read at OD450nm. The EC_90_ or EC_70_ value of viruses and the EC_50_ value of NA proteins were determined by fitting a curve with non-linear regression in GraphPad Prism and identifying the value which give 90% or 70% or 50% of NA activity. These values were used for NA inhibition assay.

For the determination of NA inhibition activity of serum or monoclonal antibodies, viruses or proteins were diluted based on EC90 or EC50 value respectively, as determined above. Monoclonal antibodies were prepared by making a serial 3-fold dilutions using assay buffer (starting from 100μg/ml). RDE-treated serum samples were prepared by making a serial of 2-fold dilutions using assay buffer (starting from 1:10 dilution). Then, 25μl of diluted monoclonal antibodies or sera was added into Fetuin-coated Maxisorp plates. Next, 25μl of diluted viruses or proteins were added and incubated at 37°C for 16-18 hours. Wells with no viruses or no rNA proteins were used as negative controls. Wells with not serum or mAbs were used as positive controls. Then, 100μl of PNA-HRP was added and incubated for 2 hours. Finally, 100μl of TMB (Sigma) substrate was added for 3 minutes before being stopped by 50μl of 0.16M sulphuric acid. For each step, plates were washed 6 times with PBS-T. Plates were read at OD450nm. NA inhibition (NI) value was calculated as followed:

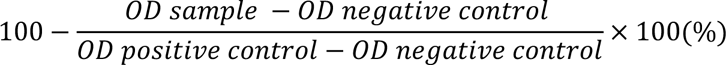

An IC_50_ value was determined by interpolating the value of serum dilution that correspond to NI of 50% using GraphPad Prism.

### MUNANA assay for NA activity

2′-(4-Methylumbelliferyl)-α-D-N-acetylneuraminic acid sodium salt hydrate or MUNANA substrate (Biosynth Carbosynth) was diluted in assay buffer (33 mM MES, 4 mM CaCl_2_, pH 6.5 in water) at final concentration of 100μM. Recombinant NA proteins were prepared by making serial 3-fold dilutions using assay buffer (starting from 10μg/ml). Next, 20μl of diluted NA proteins followed by 30μl of MUNANA were added into 96-well maxisorp black microwell plate and incubated at 37°C for 1 hour. Finally, 150μl of stop solution (0.1 M glycine, 25% Ethanol, pH 10.7 in water) was added and fluorescence intensity was read with excitation wavelength at 355nm and emission wavelength 460nm. The graph of relative fluorescence unit (RFU) corresponding to concentration of rNA was determined by fitting a curve with non-linear regression in GraphPad Prism.

### ELISA

96-well Maxisorp plates (Thermo Fisher) were coated overnight at 4°C with 2 μg/ml recombinant HA or NA proteins. After blocking with 200μl of 1% FCS in phosphate-buffered saline (PBS), duplicate wells of serially diluted sera or monoclonal antibodies were added and incubated for 2 h at room temperature. Monoclonal antibodies were prepared by making a serial 4-fold dilutions starting from 10μg/ml. Sera samples were prepared by making a serial of 4-fold dilutions starting from 1:100 dilution. Plates were washed in PBS-T (0.05% Tween- 20 in PBS) and PBS before incubation with either 1:20,000 dilution of HRP-conjugated rabbit anti-human IgG (Dako) or 1:10,0000 dilution of HRP-conjugated goat anti-mouse IgG (Seracare) for 1 h at room temperature. Plates were washed and developed using TMB substrate (Sigma) for 3 minutes, stopped by 50ul 0.16M sulphuric acid and read at 450 nm. The graph of OD corresponding to concentration of rNA was determined by fitting a curve with non-linear regression in GraphPad Prism.

### Capture ELISA

96-well Maxisorp plates (Thermo Fisher) were coated overnight at 4°C with 2 μg/ml anti-stem- HA monoclonal antibodies (CR9114) [44]. Ether-split viruses, 0.5% Tween-20-treated viruses, and 0.5% Triton-X 100-treated viruses were prepared by making a serial of 3-fold dilutions starting from 1:2 dilution. After blocking with 200μl of 1% FCS in phosphate-buffered saline (PBS), duplicate wells of serially diluted viruses were added and incubated for 2 h at room temperature. Plates were washed in PBS-T (0.05% Tween-20 in PBS) and PBS before incubation with either biotinylated anti-NA antibodies (2E01) or anti-HA antibodies (CR8033 for B/Lee/1940 virus and CR8071 for B/GL/1739/1954 virus) at concentrations of 1 μg/ml. Anti-NA antibodies (2E01) or anti-HA antibodies (CR8033 and CR8071) were previously biotinylated using biotin conjugation kit (Abcam, Fast, Type A, Lightning-Link®). Plates were washed again in PBS-T (0.05% Tween-20 in PBS) and PBS before incubation with 1:20,000 dilution of HRP-conjugated Streptavidin (Thermo scientific) for 1 h at room temperature. Plates were washed and developed using TMB substrate (Sigma) for 3 minutes, stopped by 50ul 0.16M sulphuric acid and read at 450 nm. The graph of OD corresponding to dilution of virus was determined by fitting a curve with non-linear regression in GraphPad Prism.

### Sequence analysis and antigenic cartography

Sequences were aligned using MAFFT v7.490 [45] integrated within Geneious Prime. Maximum likelihood trees incorporating the best-fit model of nucleotide substitution estimated using Smart Model Selection [46], and aBayes [47], node support were estimated in PhyML v3.0 [48]. For antigenic cartography, HI endpoint titres or NI IC_50_ values were used. Antigenic maps were generated in R using the Racmacs package [49]. Antigenic maps were constructed using 5,000 optimizations, with no minimum column basis set.

### Statistical analysis

Raw data were analysed as detailed in each section. Data were tested for normality using the Anderson-Darling test in Prism 9 (GraphPad). Correlation was assessed by computing nonparametric Spearman correlation in GraphPad Prism.

---

**Supplementary figure 1.**
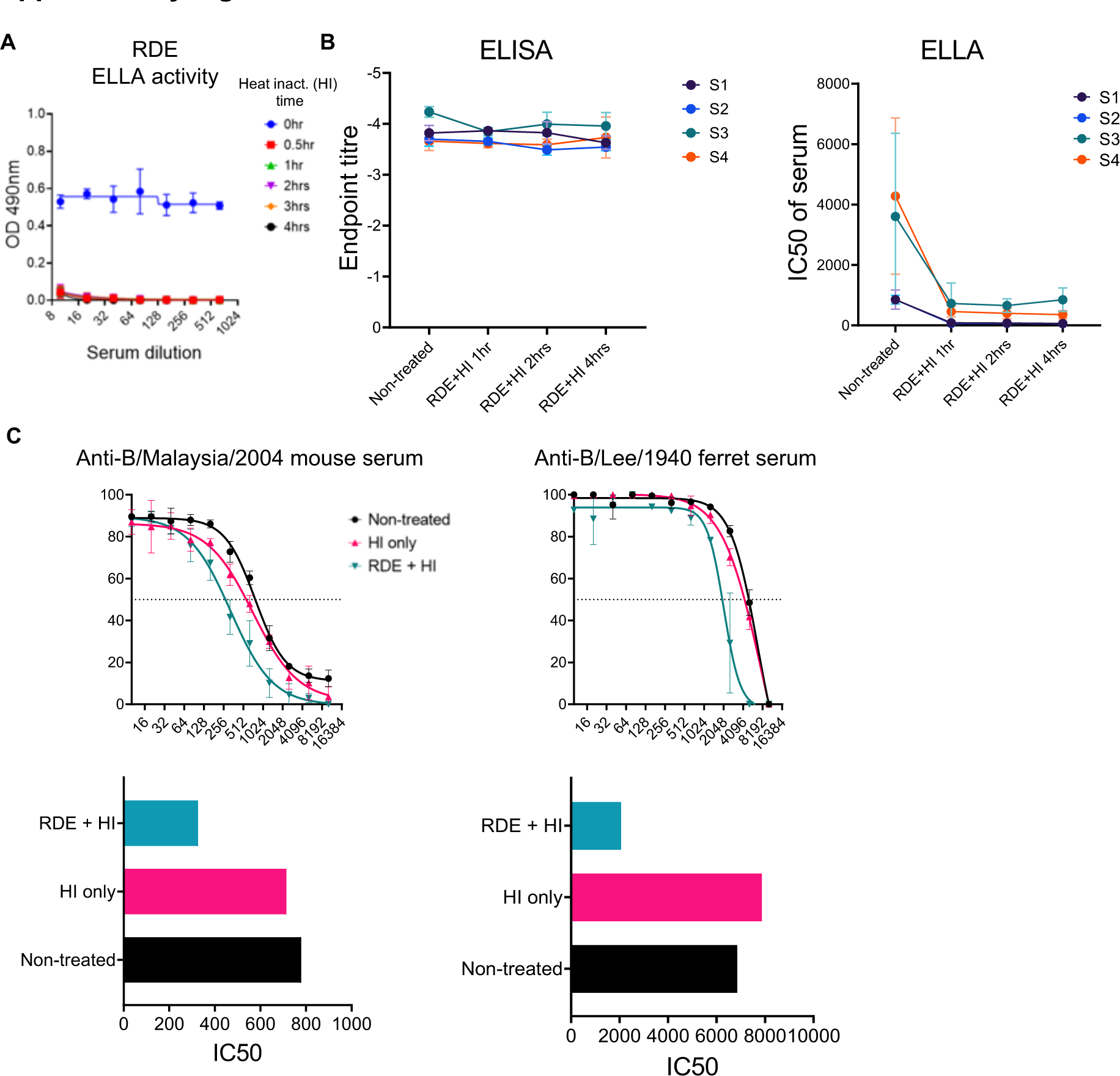
(A) Human sera (baseline) were treated with RDE and heat- inactivated for different times (0, 0.5, 1, 2, 3, or 4 hours). Sera were tested for residual RDE activity after treatment by ELLA. The mean and standard deviation are shown (n=3 donors, tested in duplicate). **(B)** Human sera (baseline, day 7, and day 28 post-vaccination) were used to check the effects of heat-inactivation on antibodies in sera by ELISA and ELLA against B/Phuket/3073/2013 rBNA. Sera were left untreated as negative controls. The mean and standard deviation are shown (n=4 donors). **(C)** Anti-B/Malaysia/2004 mouse serum and anti- B/Lee/1940 ferret serum were treated under different conditions (non-treated, heat-inactivated only, or RDE-treated and heat-inactivated) and tested by ELLA against homologous viruses. The mean and standard deviation are shown (n=2, tested in duplicate).

**Supplementary figure 2.**
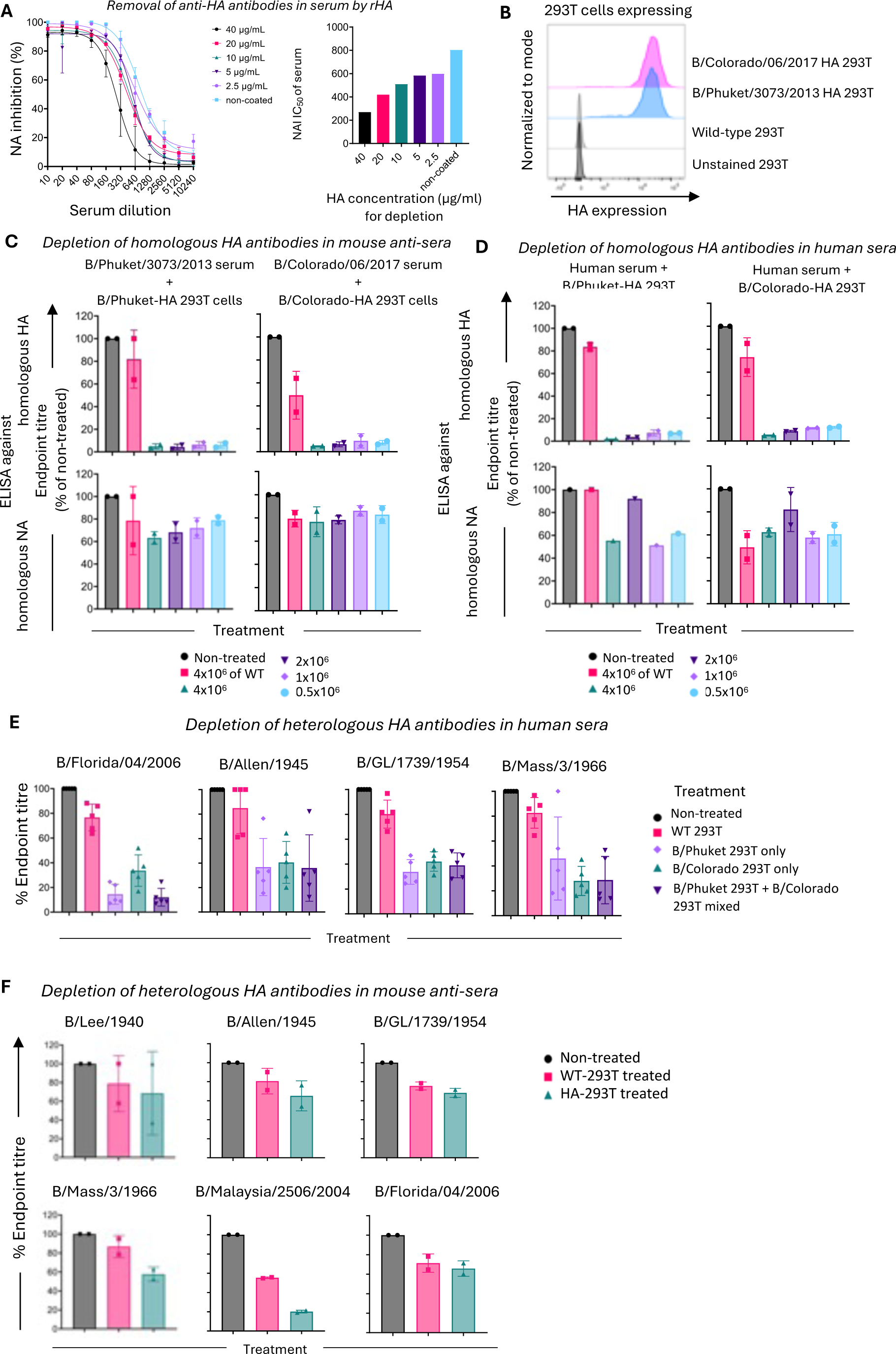
Efficiency of depleting homologous but not heterologous anti-HA antibodies in animal and human sera. (A) Anti-B/Lee/1940-HA serum was incubated on wells coated with different amounts of B/Lee/1940-HA protein prior to testing in ELLA. A non-coated well was used as negative control. **(B)** B/Phuket/3073/2013-HA and B/Colorado/06/2017-HA expressing 293T cells were stained with CR8071 and 46BH anti-HA antibodies respectively to determine levels of HA **(C)** Mouse anti-sera were either incubated with different number of HA-293T cells (ranging from 0.5x10^6^ – 4x10^6^ cells), or 4x10^6^ of WT 293T cells or left non-treated. Depletion was determined by ELISA against homologous HA and NA as a control. **(D)** Human serum samples were incubated with different number of HA-expressing 293T cells (ranging from 0.5x10^6^ – 4x10^6^ cells), or with 4x10^6^ of WT 293T cells or left non-treated. Depletion was determined by ELISA against homologous HA and NA as a control. **(E)** Human serum samples were incubated with either 4x10^6^ of B/Phuket-HA 293T cells or 4x10^6^ of B/Colorado-HA 293T cells or 4x10^6^ of a mix of two HA-expressing 293T cell lines. Sera samples were either treated with 4x10^6^ of WT 293T cells or left non-treated as negative controls. Depletion was determined by ELISA against HA from different IBV isolates. **(F)** Mouse anti-sera were incubated with 4x10^6^ of a 1:1 mixture of B/Phuket/2013-HA 293T cells and B/Colorado/2017-HA 293T cells, or with 4x10^6^ of WT 293T cells or left non-treated as negative controls. Depletion was determined by ELISA against HA of IBV strains specific for antiserum. Throughout the figure each condition was tested in duplicate, with the mean and standard deviation are shown.

